# Limb Lengthening Induces Substantial Muscle Remodeling in Humans

**DOI:** 10.64898/2026.09.27.754794

**Authors:** Hui Tang, Josh R. Baxter, Chirstine M. Goodbody, John A. Heydemann, Sarah E. Little-Letsinger, Niamh McMahon, Owen N. Beck

## Abstract

To correct lower limb length discrepancies, orthopedic surgeons often perform limb lengthening procedures. These procedures involve directly lengthening a primary bone (*e.g.,* femur), which also stretches the muscles and tendons spanning the elongation site. Limb lengthening procedures improve lower-limb length symmetry, often at the expense of chronic joint-level decrements. For example, knee joint range of motion and strength does not always return to pre-surgery values following femur lengthening. These joint-level decrements governed by altered muscle force-length characteristics, but it is unestablished how human muscles remodel during surgical limb lengthening. Here, we characterized bilateral human vastus lateralis fascicle force-length profiles following femur lengthening. At 2.2 ± 1.7 years (Avg ± SE) post-initial surgery, the lengthened limb’s vastus lateralis fascicles were longer, effectively stiffer, and weaker than the contralateral limb. The lengthened limb’s vastus lateralis fascicles were 38 mm longer (Avg) than that of the contralateral limb, even though the lengthened limb’s femur was only 31 mm longer (Avg). The onset of vastus lateralis fascicle passive force production began at 22% shorter (Avg) operating lengths in lengthened versus contralateral limb muscles. Additionally, vastus lateralis peak active force production was 34% less (Avg) in the lengthened versus contralateral limbs, and in-series quadriceps tendon stiffness was not stiffer in lengthened limbs. These data suggest that active muscle tissues lengthen more than passive muscle tissues following limb lengthening, effectively creating stiffer muscles. Years after limb lengthening, patient muscles exhibit numerous differences in their force-length properties versus that of natural growth, likely contributing to impaired joint-function.

**New & Noteworthy:** Consistent with findings in the non-human animal literature, artificially lengthened limbs house muscles that are longer, weaker, and effectively stiffer than that of the contralateral limb in humans. Here, we provide the first-in human muscle force-length data following limb-lengthening procedures, which may serve as an initial reference for healthcare professionals and help refine surgical procedures moving forward.

## 1. Introduction

Lower-limb length discrepancies are prevalent and can be present at birth or developed at any stage of life. Approximately 40-70% of people have a lower limb that is measurably longer than the other (1, 2). These estimates pertain to otherwise healthy people while also including cohorts with a relatively high prevalence of limb length discrepancy, such as those with cerebral palsy (3), skeletal dysplasia (4–6), and individuals recovering from a traumatic injury (7) or surgery (*e.g.,* hip replacement) (8, 9).

Limb-length discrepancies 2 cm cause harmful movement asymmetries during activities of daily living (10–15) and thus are routinely resolved via orthopedic limb-lengthening procedures (16). The distraction protocol involves osteotomy and an artificial device surgically affixed to the osteotomized bone. After a brief latency period, the device lengthens the bone until target segment length is achieved (17, 18). Once the distracted limb reaches target length, the bone(s) consolidates, and the lengthening device is surgically removed from the body. Throughout this multi-month procedure, patients regularly undergo physical therapy to mitigate the loss of and subsequently to regain joint function (*e.g.,* range of motion and strength) (19–25). Limb-lengthening procedures are generally satisfactory at elongating bones and inducing more symmetric whole-body locomotor patterns (26), but these procedures are plagued by high complication rates (27–35) and long-term decrements in patient joint function (36, 37).

Limb lengthening procedures regularly yield joint-level complications and functional deficits. Roughly 50% of limb lengthening complications are related to issues about joints bookending the lengthened limb segment (*e.g.,* contractures, joint subluxation, axial deviation) (27, 28, 30, 32, 34, 38–40). And lengthened-limb joints experience a reduction in range of motion (19, 22, 36, 41) and strength (23, 24, 41) for years post limb-lengthening, with some participants never regaining pre-surgery baseline joint function (37). These joint-level issues likely arise from inadequate muscle remodeling (42). However, the standard of care for limb lengthening does not directly assess patient muscle health and we are unaware of serviceable muscle data in humans, aside from a femur lengthening case study reporting that vastus lateralis fascicles elongate due to sarcomerogenesis (43). Accordingly, the specific muscle inadequacies underlying poor joint outcomes from limb-lengthening procedures are unestablished, thereby precluding improvements in the distraction protocol or rehabilitation procedures that better address patient soft-tissue needs.

While data in humans are elusive, non-human animal studies provide insight regarding how limb lengthening procedures induce muscle remodeling. In non-human animals, limb lengthening elicits longer muscle-tendon units almost exclusively through muscle lengthening via sarcomeregenesis (43–47). As such, limb lengthening procedures yield disproportionately long muscles in lengthened limbs compared to natural growth (45, 47). However, limbs with relatively longer muscles typically enable a greater joint range of motion (48), which is opposite to that observed in the post-lengthened limb joints of humans (21, 22, 36, 49, 50). Perhaps the diminished ability of muscle to remodel slack length (length at which passive tension arises) or excess extracellular matrix thickening (47, 51) contribute to effectively stiffer muscles and reduced joint range of motion after limb lengthening. Disentangling active and passive contributions to muscle force production can inform the observed decrements in joint strength across the full range of motion in the limb lengthening patient.

The goal of this short report was to provide first in-human muscle active and passive force-length profiles following lower-limb lengthening procedures. To accomplish this goal, we characterized bilateral vastus lateralis fascicle active and passive force-length profiles after femur lengthening in humans (Fig. 1). We also performed exploratory analyses of in-series quadriceps tendon stiffness to further detail soft-tissue mechanics that govern patient joint function. These data provide novel muscle-level insight regarding the factors governing poor joint-level function in limb lengthening patients.

**Figure 1.**
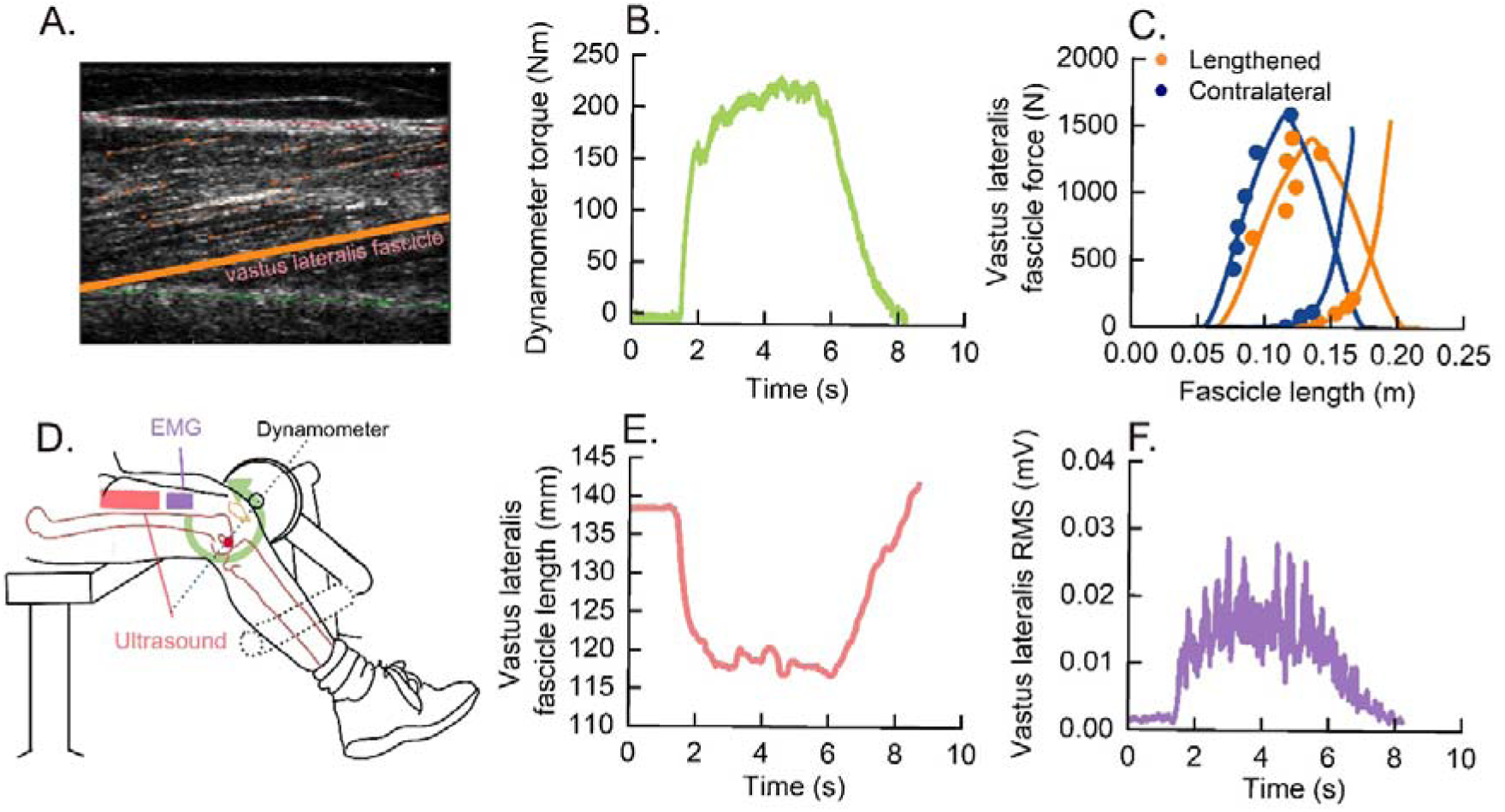
Participants performed isometric knee extension trials while researchers collected dynamometer torque, ultrasound images, and electromyography signals. A) Ultrasound image of tracked vastus lateralis fascicle (Tennessee orange line); B) dynamometer torque versus time (green); C) vastus lateralis fascicle force-length relationships for the lengthened limb (Tennessee orange) and contralateral limb (blue); D) experimental setup with a participant’s knee joint aligned with the dynamometer’s axis of rotation, ultrasound probe (pink) secured over the thigh, and a electrography sensor (purple) secured over the vastus lateralis muscle belly; E) vastus lateralis fascicle length versus time during an isometric knee extension trial (pink); and F) vastus lateralis muscle activation signal versus time (blue).

## 2. Methods

### 2.1. Participants

Three individuals who previously completed unilateral femoral lengthening using an intramedullary fixator at Central Texas Pediatric Orthopedics participated (Table 1). Each participant was free from neurological disorders. Institutional Review Boards of the University of Texas at Austin and Ascension Seton reviewed and approved the study protocol (No. 00005001). We obtained informed written assent from each participant and consent from a legal guardian before data collection.

**Table 1.** Participant Characteristics.

| Participant | Sex | Age<br>(yrs) | Height<br>(m) | Body<br>mass<br>(kg) | Pre-surgical femur length<br>(mm) |  | Post-surgical femur length<br>(mm) |  | Post-osteotomy<br>assessment<br>(week) |
| --- | --- | --- | --- | --- | --- | --- | --- | --- | --- |
|  |  |  |  |  | Lengthened | Contralateral | Lengthened | Contralateral |  |
| 1 | F | 19 | 1.82 | 77 | 478 | 486 | 506 | 485 | 183 |
| 2 | M | 14 | 1.91 | 126 | 376 | N/A | 481 | 477 | 146 |
| 3 | M | 12 | 1.53 | 49 | 373 | 383 | 391 | 387 | 10 |

### 2.2. Experimental Setup

We characterized participant bilateral vastus lateralis muscle fascicle force-length relationships, as well as bilateral quadriceps tendon force-length profiles and femur lengths. To characterize vastus lateralis muscle and tendon mechanics, we recorded dynamometer torque (100 Hz), B-mode ultrasound images (50 Hz), and muscle activation (2000 Hz), synchronized through the analog input of the motion-capture system (Vicon Nexus, Vicon Motion Systems, Oxford, UK) during active and passive trials. B-mode ultrasound images were recorded via 60 mm ultrasound probe placed over the fascicle at about halfway along the length of the thigh (Telemed, Vilnius, Lithuania) (52). We used surface electromyography (EMG) to collect muscle activity of the vastus lateralis, rectus femoris, and vastus medialis muscles during testing (Delsys Inc., Natrick, MA, 2000 Hz). We shaved and cleaned each EMG electrode site before placing each sensor superficial to the respective muscle belly, aligned with the direction of the underlying muscle fascicle. We also measured bilateral femur length from the superior end of the femoral head to the intercondylar notch from radiographs acquired before initial surgery and at the most recent post-operative follow-up assessment (Table 1; Supplementary Fig. S1).

### 2.3. Experimental Protocol

We quantified participant knee range of motion on the dynamometer setup and subsequently tested each limb at six equally spaced angles spanning the respective range of motion using the protocols described below. Notably, this range of motion was often less than that capable by the body, but was limited to the testing setup (*i.e.,* leg contacted the chair during deep knee flexion). We randomized the testing order of each knee angle per limb, and each limb per participant. At each knee angle we followed the same trial sequence. We moved the shank to the target angle, waited 45 seconds at the fixed position to allow elastic dissipative effects to settle (53), then recorded 5 seconds of relaxed dynamometer torque, muscle activation, and ultrasound images. Next, we instructed the participant to perform a maximal voluntary isometric knee extension contraction (MVC). We provided visual feedback of the dynamometer torque trace and verbal encouragement during each MVC. Between each MVC trial, participants rested at least 2 minutes between to minimize the effects of fatigue. Subsequently, we repositioned the ultrasound probe over the vastus lateralis muscle–tendon junction and recorded two ramped MVC trials at each of the middle knee joint angles. We analyzed the trial with the best image quality.

### 2.4. Data Analysis

We computed active knee extension moments. We filtered dynamometer torque (τ_dyno_) using a fourth-order Butterworth low-pass filter (10 Hz). To isolate the moment produced by the knee (*M_knee_*), we subtracted torque due to the dynamometer attachment (τ_attachment_) and the shank and foot due to gravity (τ_leg_) consistent with segment-mass regression equations (54) (Eq.1):

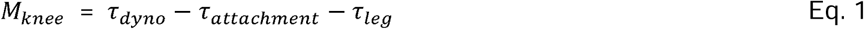

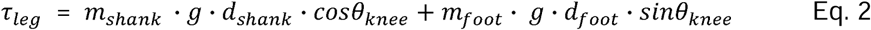

To estimate torque due to the leg (Eq. 2), we deemed shank mass (m_shank_) and foot mass (m_foot_) as 4.65% and 1.45% of body mass, respectively; the distance of the knee joint to the shank center of mass (d_shank_) as 0.433 of shank length from the knee; the distance from the knee joint to the foot center of mass (d_foot_) as shank length plus half of foot length; and we used knee angle relative to vertical (θ_knee_); and gravitational acceleration (g). We measured shank length (from the lateral femoral condyle to the lateral malleolus) and foot length (from the heel to the tip of the longest toe) using a tape measure.

We calculated vastus lateralis muscle fascicle forces at each testing trial (Eq. 3). At each testing angle, we divided knee joint moment ( by the knee joint moment arm (*r_knee_*), multiplied by the fractional contribution of the vastus lateralis to total quadriceps physiological cross-sectional area (A_VL_ = 0.34) (55), and divided by the cosine of the vastus lateralis pennation angle with respect to the aponeurosis (θ_pen_):

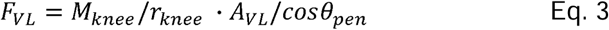

We computed the knee joint to vastus lateralis moment arm by measuring its distance at a knee angle of 90° and using a participant-specific-scaling angle-dependent regression (56).

We extracted vastus lateralis fascicle length and pennation angle for each active and passive knee extension trial from the ultrasound images using a semi-automatic tracking algorithm (52, 57). The active fascicle length during each MVC (L_active_) was computed as the mean of five consecutive ultrasound frames centered on the frame with peak knee moment. The passive fascicle length at each rest trial (L_passive_) was the mean over the 5-second relaxed segment recorded immediately before the MVC at the same knee angle.

We fit each limb’s passive force-length relationships. Passive fascicle forces and their corresponding lengths across joint angles were modeled using an exponential equation (58) (Eq. 4)

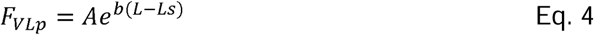

In Eq. 4, L is the fascicle length; A and b are constants determined using nonlinear least-squares fitting (lsqcurvefit function using MATLAB, MathWorks, Natick, MA, USA); and Ls is the fascicle’s slack length, the fascicle length at which the passive force first exceeds a 50 N threshold (59, 60). We set all negative raw values to zero before fitting the exponential. We verified that vastus lateralis activation remained below 10% of each participant’s maximal voluntary EMG amplitude during all passive trials (Avg ± SE: 6.5 ± 2.3%).

We computed each limb’s active fascicle force-length relationships. Force production at relatively long fascicle lengths is contributed by both active and passive elements. We therefore subtracted the passive fascicle force from the respective muscle length from the peak total force production to isolate active force contribution:

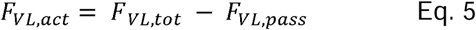

Where *F_VL,tot_* is the fascicle force computed from the peak MVC torque by Eq. 3, and *F_VL,pass_* is the passive force (Eq. 4) from the corresponding muscle fascicle length. We identified the optimal muscle fascicle length for active force production (L_0_) by minimizing the squared residuals between the experimental data and the OpenSim active force–length spline (61, 62), using fmincon (MATLAB R2024a, MathWorks, Natick, MA, USA) with L_0_ bounded [0.06 m, 0.22 m].

We quantified vastus lateralis quadriceps tendon force-displacement profiles. We calculated tendon force (*F*_T_) at 11 discrete percentages (0, 10, 20,…, 100%) of each limb’s peak muscle-tendon force during the ramped MVC trial. At each of 11 data points,*F_T_* was the total quadriceps tendon force computed by dividing the knee joint moment *M_knee_* by the knee joint moment arm (*r_knee_*)evaluated at the knee angle at which the ramped MVC was performed (Eq. 6):

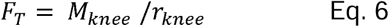

We tracked the position of the vastus lateralis muscle–tendon junction during the MVC using B-mode ultrasound, EchoWave software (Telemed, Vilnius, Lithuania), and custom MATLAB script. In-series vastus lateralis tendon displacement was quantified as the displacement of the muscle–tendon junction with respect to a zero active force reference position. We then fit a 2^nd^ order polynomial to the tendon force-displacement data points (63). For each participant, we calculated the tendon stiffness (k_t_) as the change in tendon force divided by the corresponding change in tendon displacement (*L_r_*) (Eq. 7):

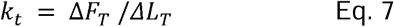

We reported all calculated results in average ± standard error (Avg ± SE).

### 2.5 Data Quality

We assessed muscle and tendon force versus length relationships data quality using multiple analyses. First, we computed intraclass correlation coefficients (ICC) (64) and coefficients of determination (R²). Next, we assessed parameter sensitivity to individual data points using a leave-one-out (LOO) procedure (65): the coefficient of variation (CV) of the resulting N estimates is reported for L₀, F_max_, and *k_t_*. We pre-specified three data-retention thresholds: active and passive force-length relationship ICC ≥ 0.7 and R² ≥ 0.5. We also reported and leave-one-out CV for the respective variables (66). All analyses used custom scripts in MATLAB (R2024a, MathWorks, Natick, MA). We interpreted data that passed the quality inspection in the results section and presented full analysis in the supplementary materials.

## 3. Results

Vastus lateralis fascicles in the lengthened limbs produced peak active force at longer lengths than that of the contralateral limb (Fig. 2). Optimal vastus lateralis fascicle length for active force production was 68 mm longer in the lengthened limb of participant 1, 20 mm longer in participant 2, and 27 mm longer in participant 3 compared to the respective contralateral limb; yielding 32% ± 25% (Avg ± SE) longer fascicles in patient lengthened versus contralateral limbs.

**Figure 2.**
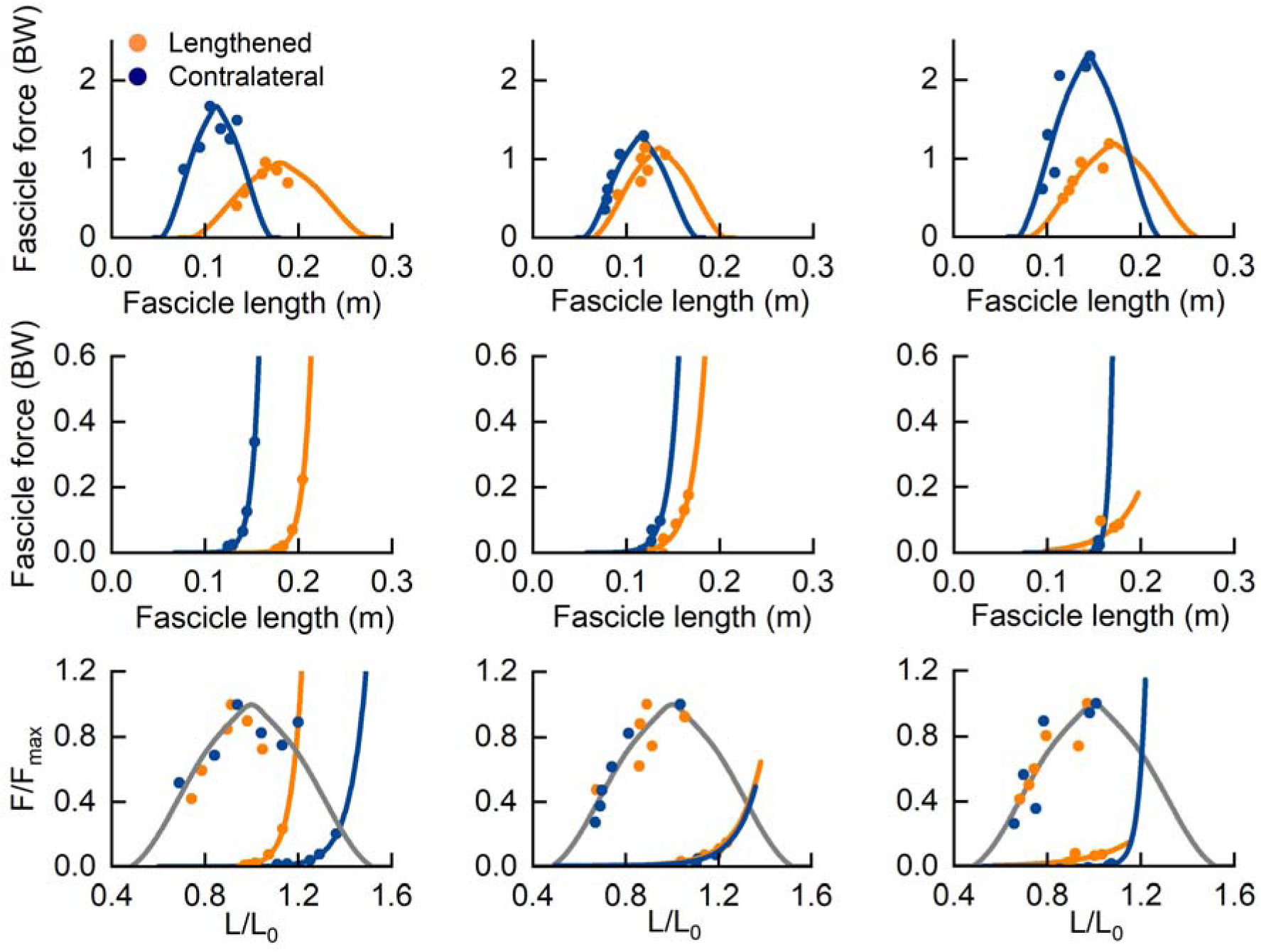
Post-limb lengthening vastus lateralis muscle force-length relationships. Each column represents one participant. Rows represent (top) active and (middle) passive muscle fascicle force in units of body weight (BW) versus length (m), as well as (bottom) muscle fascicle force as a fraction of muscle fascicle active force capacity (F/F_max_) versus length as a fraction of optimal length for active force production (L/L_0_). Tennessee orange data represent the lengthened limb and blue data represent the contralateral control limb.

Vastus lateralis fascicles in the lengthened limbs began producing passive tension at longer fascicle lengths than the contralateral limb (Fig. 2). The lengthened limb’s vastus lateralis fascicle slack length occurred at a 54 mm longer length than that of the contralateral limb in participant 1, and at a 19 mm longer length in participants 2 and 3; yielding 22 ± 14% (Avg ± SE) longer fascicle slack lengths in lengthened versus contralateral limbs. Notably, the vastus lateralis muscle fascicle slack length occurred at a relatively shorter length with respect to the optimal length for active force production in each of the lengthened limb muscles compared to the contralateral limb.

Femurs in the lengthened limbs were longer than that of the contralateral limbs post limb lengthening (Table 1). Compared to the contralateral limb, participant 1’s lengthened femur was 21 mm longer, and participant 2 and 3’s lengthened femurs were 4 mm longer in the most recent x-ray image. These inter-limb bone-length differences were smaller than those of attached muscles. Numerically, lengthened limb vastus lateralis muscle fascicles, measured using the optimal length for active fascicle force production (L_0_), were 38 mm longer (Avg) than those on the contralateral limb, whereas the underlying lengthened femur was only 31 mm longer (Avg).

The lengthened vastus lateralis produced less peak active fascicle force than the contralateral muscle in every participant (Fig. 2). Peak vastus lateralis fascicle force was 42% lower in the lengthened versus contralateral limb in participant 1, 10% lower in participant 2, and 48% lower in participant 3; yielding 33 ± 21% (Avg ± SE) weaker muscles in patient lengthened versus contralateral limb muscles.

The lengthened limb’s quadriceps tendon was not stiffer than that of the contralateral limb. Compared to the contralateral limb, average quadriceps tendon stiffness was numerically less in the lengthened limb in two participants and nearly identical in the third participant (Fig. 3). Thus, changes in tendon stiffness do not drive the decrements in patient joint range of motion.

**Figure 3.**
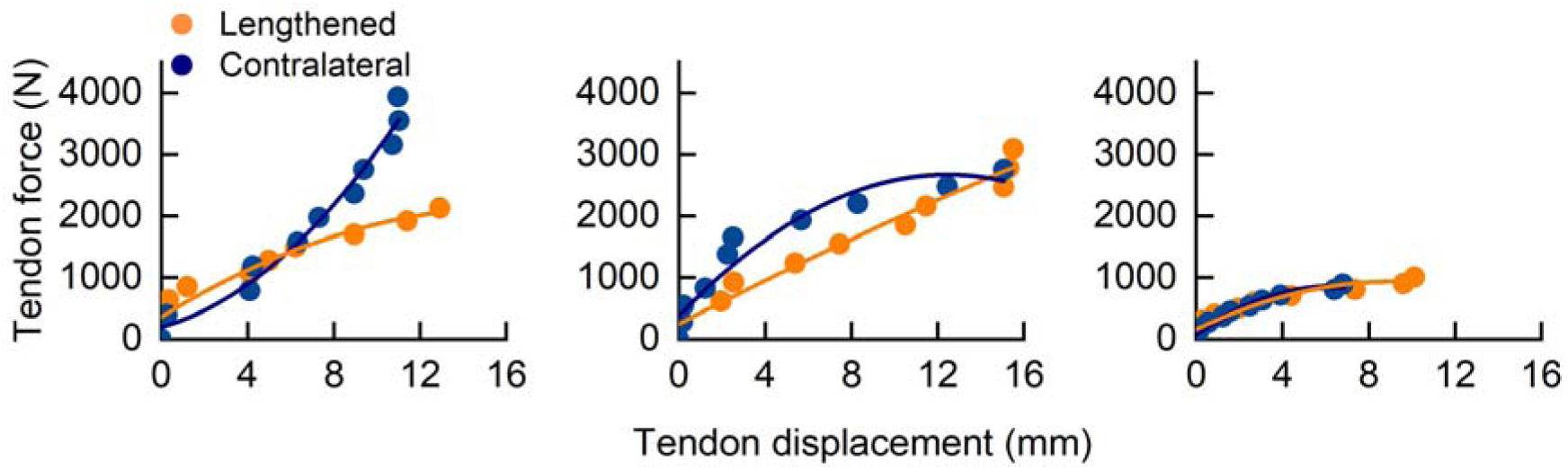
Post limb lengthening quadriceps tendon force-displacement relationships. Each column represents a participant’s tendon force versus displacement profile. Symbols are of lengthened (orange) and contralateral (blue) limbs post-distraction. Lines are force-displacement curves fitted to 2^nd^ polynomial fit, which in all cases exceeded R^2^ =0.9.

## 4. Discussion

Here, we present the first in-human muscle-tendon force-length profiles following limb lengthening procedures, which may serve as an initial reference for healthcare professionals and guide improvement of limb lengthening surgical protocols. Our results are consistent with findings in the non-human animal literature, as the lengthened limbs house muscles that are longer, effectively stiffer, and weaker than the contralateral limb. Inducing lower-limb length symmetry in humans via unilateral femoral lengthening inadvertently yields artificially long vastus lateralis fascicles in lengthened limbs. Yet, the increased length of active muscle contractile elements is not matched by passive muscle elements, yielding muscles that produce passive tension at relatively shorter operating lengths. In-series to the vastus lateralis, quadriceps tendons were not stiffer in the lengthened versus contralateral control limbs, suggesting that changes in tendon stiffness do not drive the decrements in patient joint range of motion. Further experiments that longitudinally quantify and associate changing muscle-tendon mechanics with joint function throughout limb-lengthening procedures are needed for mechanistic insight regarding patient outcomes.

Human muscles are substantially longer in lengthened limbs versus natural growth. Surgically elongating the femur, which extends the distance between the origin and insertion of the vastus lateralis muscle-tendon unit, caused muscle fascicles to lengthen more than the underlying bone. The greater lengthening of vastus lateralis muscle fascicles versus the femur due to surgical lengthening in humans has been suggested in a case study (using coarse methods) (43), and nearly matched muscle and bone lengthening has been reported in non-animal human studies (45). Previous studies also suggest that these muscle length changes are due to sarcomeregenesis (43, 45–47, 67), which enable muscles to operate closer to the plateau region of their active force-length profile during typical movements (68). While it is not surprising that muscle fascicles spanning the bone’s distraction site lengthen, the enormous effect size is curious. It is unlikely that muscles need to lengthen more than the distracted bone to maintain operating along economical regions of the force-length profile, but we did not assess muscle fascicle lengths longitudinally nor during activities of daily living to verify.

Compared to sarcomeres, the tissues that govern passive muscle force-length profiles modestly remodel their length. In our participants, the lengthened-limb vastus lateralis fascicles begin producing passive tension at 7% shorter operating lengths (L_s_/L_0_) than the contralateral control limb. Similarly, non-human animal studies report that lengthened-limb muscle slack length does not typically lengthen to the extent of the optimal muscle length for active force production, and that this length-change differential is associated with fibrosis and excess thickening of the endomysium and perimysium (47, 51). Furthermore, human joint range of motion is reduced due to limb lengthening (21, 22, 36, 49, 50) and overall muscle-tendon units from non-human animals are 15% stiffer in the lengthened versus control limbs (69). Being that tendons do not increase stiffness during limb lengthening (Fig. 3) (70), it is likely the muscles that stiffen during limb lengthening procedures. Functionally, muscles that develop passive force at relatively shorter lengths have a diminished ability to operate across the force-length profile range (71, 72), have a reduced active muscle force capacity (73), and operate at less economical lengths during walking (72, 74). Therefore, the consideration of passive muscle remodeling during limb lengthening procedures may be a worthwhile endeavor.

The present report’s novel assessment of limb-lengthening patient muscle-tendon mechanics is not without potential limitations. We caution over-interpreting the results due to modest sample size and cross-sectional observation. The absence of pre-limb lengthening muscle-tendon data prevents us from definitively revealing how soft tissues remodel due to limb lengthening, but the cross-sectional comparisons versus the contralateral limb are directionally consistent with longitudinal studies involving non-human animals. Additionally, our non-invasive measurement technique was practical for studying pediatric limb-lengthening patients but involves more assumptions than invasive protocols. For example, our 2D ultrasound images cannot account for out-of-plane movement, and our muscle force estimates depend on assumptions of force sharing, consistent neural drive across testing angles, and minimal co-activation. These and other assumptions are generally acceptable for studies of vastus lateralis force-length profiling (52, 55, 75), and we implemented secondary assessments to several critical assumptions (*e.g.,* lower-limb muscle activation was <10% of the maximum during passive trials).

Limb-lengthening procedures benefit patients by improving lower-limb symmetry, often at the expense of chronic joint-level decrements (49, 76). We posit that these joint-level decrements primarily stem from inadequate muscle remodeling. In this short report, we characterized bilateral muscle-tendon force-length profiles after femur lengthening procedures to inform healthcare professionals of patient soft-tissue health. These data may be useful for informing clinical decisions (*e.g.,* selecting specific physical therapy exercises) and motivating clinical trials with the long-term goal of improving limb-lengthening patient outcomes.

## Supporting information

Supplementary Results

## Acknowledgement

No sponsor to be declared by all authors.

## Author Contributions

O.N.B conceived and designed the research; H.T., N.M., & J.A.H. performed experiments; H.T. analyzed data; H.T., J.A.H., J.R.B, and O.N.B., interpreted results of experiments; H.T. and O.N.B. prepared figures; H.T. and O.N.B. drafted manuscript; H.T., J.A.H., J.R.B., N.M., S.E.L., and O.N.B. edited and revised manuscript; H.T., J.R.B., J.A.H., N.M., C.M.G., S.E.L., and O.N.B. approved final version of manuscript.

## Supplementary Material

Supplementary results include data quality summary, radiographic measurements (Table S1& Figure S1), muscle activation (Figure S2), vastus lateralis muscle-tendon force-length profiles data quality (Table S2, S3, S4, and S5).

## Data Availability

Data for this study are available at https://doi.org/10.6084/m9.figshare.33334533

