## Supplementary Results for "Limb Lengthening Induces Substantial Muscle Remodeling in Humans"

**Short Report**

**Data quality**

Five of six limbs passed the pre-specified fit-quality thresholds for both active and passive analyses, and every limb passed for the tendon analysis. For the passive vastus lateralis fascicle force-length fits, ICC ranged from 0.72 to 0.99 and R² from 0.55 to 0.98 (Supplementary Results Table S2). Five of six limbs exceeded the ICC ≥ 0.75 threshold; Participant 3’s lengthened limb fell short (ICC = 0.72, R² = 0.55), reflecting scattered passive-tension observations on that limb. For the active fascicle force-length fits, ICC ranged from 0.79 to 0.97 and R² from 0.41 to 0.93 (Supplementary Results Table S3). All six limbs met the ICC ≥ 0.75 threshold; Participant 1’s contralateral limb fell short (R² = 0.41), reflecting a data cluster that lay entirely on the ascending limb and therefore constrained the OpenSim spline shape only weakly; the within-participant lengthened-versus-contralateral L₀ direction was preserved when the single most influential MVC was removed. For the tendon force–displacement polynomial fits, ICC ranged from 0.97 to 0.99 and R² from 0.93 to 0.98 (Supplementary Results Table S4); all six limbs exceeded the thresholds.

Leave-one-out sensitivity was within the 5% pre-specified threshold for L_0_ and L_slack_ in every limb but exceeded 5% for F_max_ in two limbs (Supplementary Results Table S5). L_0_ leave-one-out CV ranged from 1.54% to 4.14% and L_slack_ from 0.17% to 1.41%. F_max_ leave-one-out CV ranged from 2.36% to 8.36%, exceeding 5% in the lengthened limb of Participant 3 (8.36%) and the contralateral limb of Participant 2 (7.49%). This elevated F_max_ sensitivity reflects the leverage of the single highest-force MVC on the observed maximum force rather than a deficiency in the underlying data; the direction of every lengthened-versus-contralateral F_max_ difference was preserved when the most influential MVC was removed. All six limbs were therefore retained in the primary analysis.

**Bone lengths:**

We collected three participants’ X-ray examined limb lengths before and after surgery (Figure S1). With two experienced pediatric orthopedical physicians’ assistance, we measured bilateral total limb lengths (from the most superior aspect of the femoral head to the tibial plafond) and femur lengths (from the most superior aspect of the Great trochanter to the center of femoral notch) from X-ray images. We acknowledged that measurement errors might come from participants’ standing postures, especially with ortho blocks under the shorter limb; as well as other lower limb issues such as mechanical axis misalignments.

**Table S1. Three participants’ X-ray examined bone lengths.**

| Participant | Condition | Examination day  (day) | Total limb length (mm) | | Femur length (mm) | |
| --- | --- | --- | --- | --- | --- | --- |
|  |  |  | Lengthened | Contralateral | Lengthened | Contralateral |
| 1 | Pre-surgery | 74 before surgery | 901.65 | 935.58 | 477.56 | 486.33 |
|  | Post-surgery | 1198 after surgery | 931.64 | 931.58 | 505.89 | 484.6 |
| 2 | Pre-surgery | 870 before surgery | 718.81 | 765.27 | 375.62 | Missing |
|  | Post-surgery | 922 after surgery | 956.86 | 958.31 | 481.34 | 477.22 |
| 3 | Pre-surgery | 60 before surgery | 717.14 | 739.36 | 373.19 | 382.84 |
|  | Post-surgery | 100 after surgery | 739.31 | 739.32 | 390.72 | 386.91 |

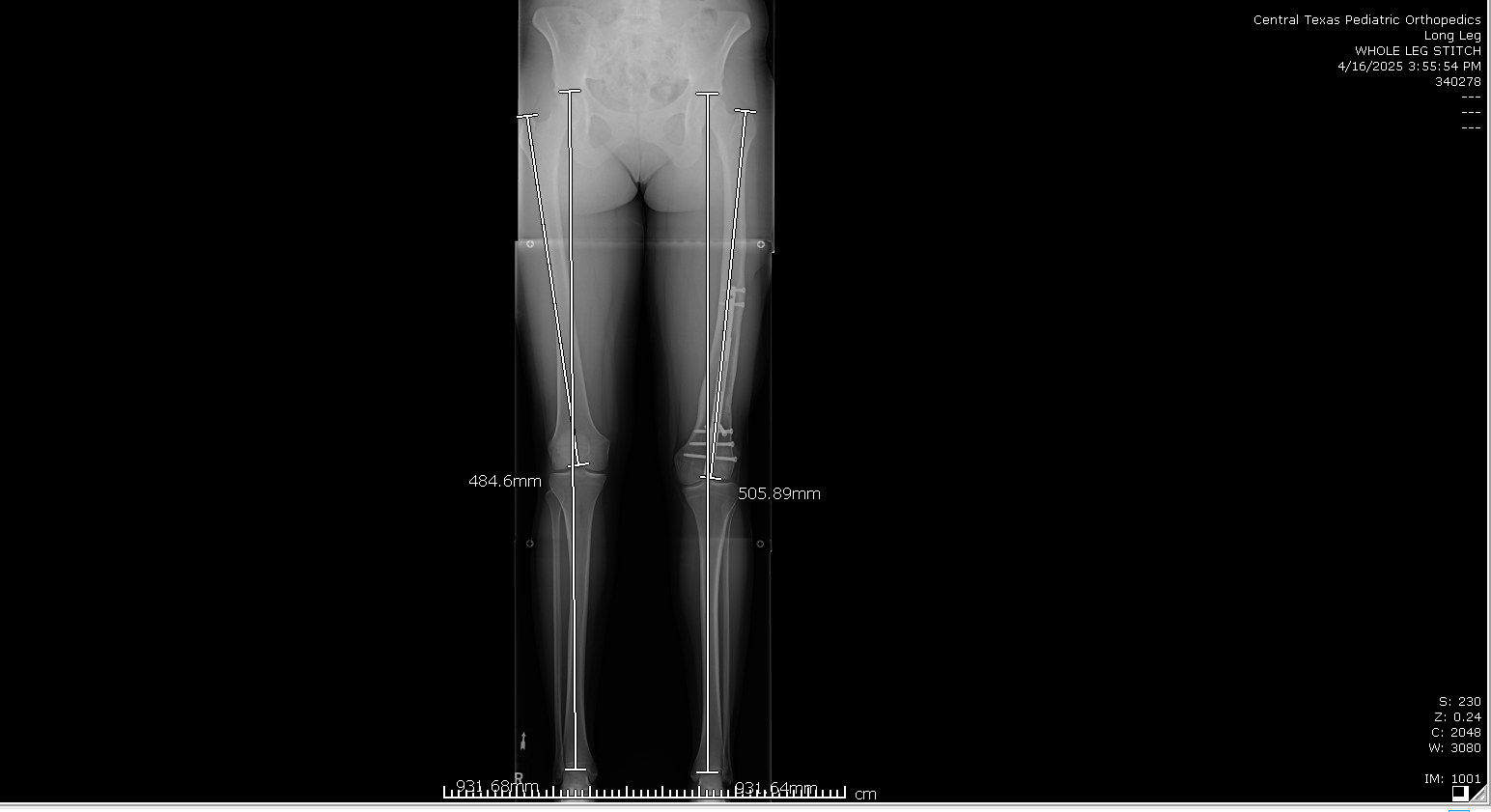

**Figure S1.** Representative standing anteroposterior radiograph illustrating the total limb length and femur length measurement used throughout this study. Femur length was measured bilaterally from the superior aspect of the femoral head to the intercondylar notch, indicated by the annotated landmarks.

**Muscle activation:** We verified vastus lateralis, vastus medialis and rectus femoris muscle activation during passive trials (Fig. S1). During passive trials across different joint angles, three participants’ quadriceps muscle activation was lower than 10% of MVC’s muscle activation (Avg ± SE P1: vastus medius: 4.09 ± 3.73, rectus femoris: 4.63 ± 4.31, vastus lateralis: 5.44 ± 3.80; P2: vastus medius: 4.77 ± 3.31, rectus femoris: 2.64 ± 1.53, vastus lateralis: 9.37 ± 4.93; P3: vastus medius 5.08 ± 2.54, rectus femoris: 4.09 ± 5.29, vastus lateralis: 4.84 ± 7.15).

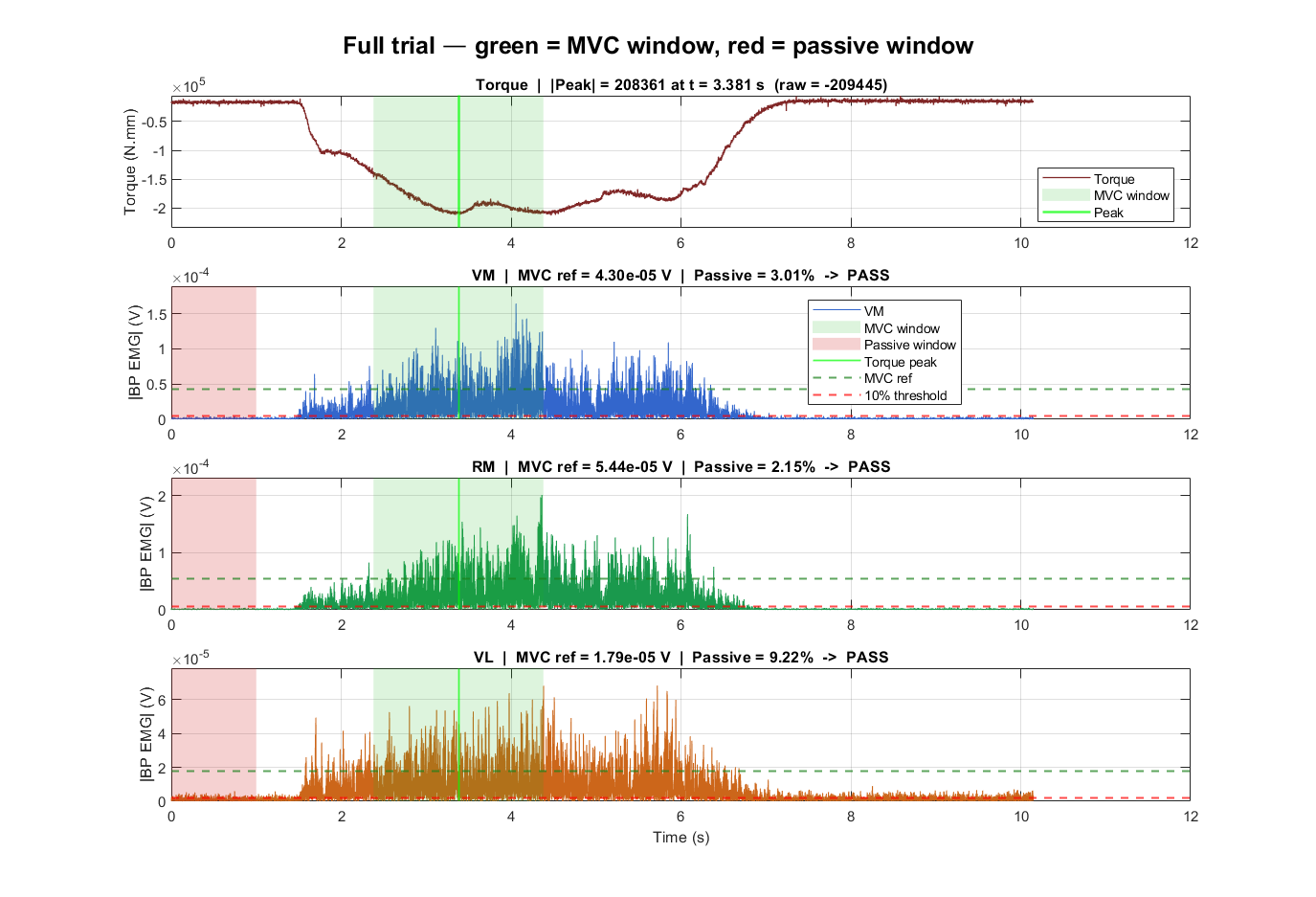

**Figure S2.** Representative muscle activation of quadricep muscles during maximal voluntary contraction. From the the top to bottom panels, there were participant’s MVC torque output, vastus medius, rectus femoris, and vastus lateralis muscle activation profiles, respectively. We matched the passive trial window as well as the MVC peak window in color red and green shades respectively and the peak MVC was labeled in light green.

**Vastus lateralis muscle/tendon force-length/displacement profiles data quality:** We reported the complete fit-quality and sensitivity statistics for every vastus lateralis passive force–length curve, active force–length curve, and in-series tendon force–displacement curve analyzed in the main manuscript, together with the per-observation data underlying each fit. Six limbs (three participants × two limbs) are reported throughout. Values that failed a pre-specified retention threshold are marked with an asterisk and identified in the accompanying footnote.

### Supplementary Table S2. Passive fascicle force–length fit parameters and quality

| **Participant** | **Limb** | **L**_slack_ **(mm)** | **ICC** | **R²** |
| --- | --- | --- | --- | --- |
| 1 | Lengthened | 193.9 | 0.999 | 0.998 |
|  | Contralateral | 140.0 | 0.999 | 0.997 |
| 2 | Lengthened | 145.9 | 0.978 | 0.949 |
|  | Contralateral | 126.7 | 0.950 | 0.895 |
| 3 | Lengthened | 180.7 | 0.724* | 0.545 |
|  | Contralateral | 161.8 | 0.913 | 0.818 |

Passive fascicle force was fit with F = A · exp(b(L − Ls)) (Eq. 4, main text). L_slack_ is the operational slack length, the fascicle length at which the fitted passive force first exceeds 50 N. ICC is the two-way random-effects, absolute-agreement, single-measure intraclass correlation coefficient between observed and fitted passive force. * denotes a value below the pre-specified retention threshold (ICC ≥ 0.75; R² ≥ 0.5). Only the lengthened limb of Participant 3 fell below a threshold, failing on ICC (0.724) while marginally satisfying R² (0.545). Its passive prediction was still subtracted from the measured MVC before the active fit, but its passive profile is not interpreted in the main text*.*

### Supplementary Table S3. Active fascicle force–length fit parameters and quality

| **Participant** | **Limb** | **L**_0_ **(mm)** | **F**_max_ **(N)** | **F_max_ (BW)** | **ICC** | **R²** |
| --- | --- | --- | --- | --- | --- | --- |
| 1 | Lengthened | 180.3 | 720.6 | 9.4 | 0.786 | 0.562 |
|  | Contralateral | 112.2 | 1258.7 | 16.3 | 0.785 | 0.414* |
| 2 | Lengthened | 135.0 | 1411.4 | 11.2 | 0.805 | 0.544 |
|  | Contralateral | 115.1 | 1589.4 | 12.4 | 0.965 | 0.930 |
| 3 | Lengthened | 171.3 | 571.3 | 11.7 | 0.916 | 0.788 |
|  | Contralateral | 144.4 | 1107.1 | 22.6 | 0.879 | 0.754 |

* denotes a value below the pre-specified retention threshold (ICC ≥ 0.75; R² ≥ 0.5). The contralateral limb of Participant 1 fell below the R² threshold because all six observations lay on the ascending limb of the force–length curve, weakly constraining the spline shape.

### Supplementary Table S4. In-series vastus lateralis tendon force–displacement fit quality and stiffness

| **Participant** | **Limb** | **Peak F**_MTU_ **(N)** | **F**_peak,weaker_ **(N)** | **ΔL at F**_peak,weaker_ **(mm)** | **k (N/mm)** | **ICC** | **R²** | **K LOO CV (%)** |
| --- | --- | --- | --- | --- | --- | --- | --- | --- |
| 1 | Lengthened (weaker) | 2132.7 | 2132.7 | 13.52 | 157.8 | 0.97 | 0.93 | 2.4 |
|  | Contralateral | 3943.5 | 2132.7 | 8.01 | 266.3 | 0.99 | 0.98 | 0.8 |
| 2 | Lengthened | 3090.9 | 2757.2 | 14.95 | 184.4 | 0.99 | 0.97 | 2.8 |
|  | Contralateral (weaker) | 2757.2 | 2757.2 | 15.14 | 182.1 | 0.97 | 0.93 | 1.2 |
| 3 | Lengthened | 1003.5 | 895.7 | 8.48 | 105.6 | 0.97 | 0.93 | 3.6 |
|  | Contralateral (weaker) | 895.7 | 895.7 | 7.04 | 127.3 | 0.99 | 0.98 | 2.5 |

Tendon force–displacement data were fit with a second-order polynomial through 11 discrete force levels per limb (Section 3.4, main text). Fpeak,weaker is the smaller of the two limbs’ peak in-series tendon force within each participant and defines the common force at which stiffness was evaluated for both limbs of that participant. K = Fpeak,weaker / ΔL(Fpeak,weaker). All six limbs exceeded both retention thresholds. K values are comparable within but not across participants, because each participant defines their own Fpeak,weaker.

### Supplementary Table S5. Leave-one-out sensitivity of fitted parameters

| **Participant** | **Limb** | **L**_0_ **CV (%)** | **L**_0_ **max-Δ (%)** | **F**_max_ **CV (%)** | **F**_max_ **max-Δ (%)** | **L**_slack_ **CV (%)** | **L**_slack_ **max-Δ (%)** |
| --- | --- | --- | --- | --- | --- | --- | --- |
| 1 | Lengthened | 2.93 | 5.36 | 4.20 | 10.11 | 0.22 | 0.39 |
|  | Contralateral | 2.16 | 3.61 | 4.48 | 10.78 | 0.17 | 0.30 |
| 2 | Lengthened | 1.54 | 2.32 | 3.22 | 7.78 | 1.33 | 2.16 |
|  | Contralateral | 2.54 | 5.79 | 7.49* | 17.81 | 1.01 | 2.08 |
| 3 | Lengthened | 4.14 | 9.70 | 8.36* | 19.81 | 1.36 | 2.26 |
|  | Contralateral | 2.82 | 4.10 | 2.36 | 5.73 | 1.41 | 3.04 |

Leave-one-out analysis removed one observation at a time and refit the corresponding curve, yielding N estimates of each parameter per limb (N = 6). CV is the coefficient of variation of those N estimates; max-Δ is the largest percentage deviation of any single leave-one-out estimate from the full-data value. The pre-specified retention threshold was CV ≤ 5% for L_0_. * denotes a value exceeding 5%. L_0_ met the threshold in every limb (1.54–4.14%) as did Lslack (0.17–1.41%). Fmax exceeded 5% in two limbs, reflecting the leverage of the single highest-force MVC on the observed maximum; the direction of every lengthened-versus-contralateral Fmax difference was preserved when the most influential MVC was removed.
